# Cell-Type-Specific Decrease of the Intrinsic Excitability of Motor Cortical Pyramidal Neurons in Parkinsonism

**DOI:** 10.1101/2020.10.20.347732

**Authors:** Liqiang Chen, Samuel Daniels, Yerim Kim, Hong-Yuan Chu

## Abstract

The hypokinetic motor symptoms of Parkinson’s disease (PD) are closely linked with a decreased motor cortical output as a consequence of elevated basal ganglia inhibition. However, whether and how the loss of dopamine alters the cellular properties of motor cortical neurons in PD remains undefined. We induced parkinsonism in adult C57BL6 mice of both sexes by injecting neurotoxin, 6-hydroxydopamine, into the medial forebrain bundle. By using *ex vivo* patch-clamp recording and retrograde tracing approach, we found that the intrinsic excitability of pyramidal tract neurons (PTNs) in the motor cortical layer 5b was greatly decreased in parkinsonism; but the intratelencephalic neurons (ITNs) were not affected. The cell-type-specific intrinsic adaptations were associated with a depolarized threshold and broadened width of action potentials in PTNs. Moreover, the loss of midbrain dopaminergic neurons impaired the capability of M1 PTNs to sustain high-frequency firing, which could underlie their abnormal pattern of activity in the parkinsonian state. We also showed that the decreased excitability in parkinsonism was caused by an impaired function of both persistent sodium channels and the large conductance, Ca^2+^-activated K^+^ channels. Acute activation of dopaminergic receptors failed to rescue the impaired intrinsic excitability of M1 PTNs in parkinsonian mice. Altogether, our data demonstrated a cell-type-specific decrease of the excitability of M1 pyramidal neurons in parkinsonism. Thus, intrinsic adaptations in the motor cortex, together with pathological basal ganglia inhibition, underlie the decreased motor cortical output in parkinsonian state and exacerbate parkinsonian motor deficits.

**Significance statement:** The degeneration of midbrain dopaminergic neurons in Parkinson’s disease remodels the connectivity and function of cortico–basal ganglia–thalamocortical network. However, whether and how dopaminergic degeneration and the associated basal ganglia dysfunction alter motor cortical circuitry remain undefined. We found that pyramidal neurons in the layer 5b of the primary motor cortex (M1) exhibit distinct adaptations in response to the loss of midbrain dopaminergic neurons, depending on their long-range projections. Besides the decreased thalamocortical synaptic excitation as proposed by the classical model of Parkinson’s pathophysiology, these results, for the first time, show novel cellular and molecular mechanisms underlying the abnormal motor cortical output in parkinsonism.

## Introduction

The degeneration of dopamine (DA) neurons in the substantia nigra (SN) alters the connection and computation of cortico–basal ganglia–thalamocortical network, which underlies the devastating motor symptoms in Parkinson’s disease (PD), including akinesia, bradykinesia, and rigidity (Albin et al., 1989; Galvan and Wichmann, 2008; McGregor and Nelson, 2019). Specially, the loss of SN DA neurons increases and decreases the activities of indirect and direct pathways, respectively, which disrupts the balanced activity between striatal direct and indirect pathways, leading to the hypokinetic symptoms in PD (Albin et al., 1989; DeLong, 1990). The pathway-specific alterations in the striatum following the loss of SN DA neurons induce numerous cellular and synaptic changes in the basal ganglia nuclei and extended brain regions, which further drive the abnormal neural activity throughout the cortico–basal ganglia– thalamocortical network in parkinsonism (Gittis et al., 2011; Kita and Kita, 2011; Fieblinger et al., 2014; Chu et al., 2015, 2017; Mathai et al., 2015; Shen et al., 2015; Parker et al., 2016, 2018; Sharott et al., 2017; McIver et al., 2019; Willard et al., 2019).

The primary motor cortex (M1) plays essential and complex roles in motor control and motor learning and is a key node in the cortico–basal ganglia–thalamocortical network (Shepherd, 2013; Ebbesen and Brecht, 2017; Ebbesen et al., 2018). M1 is a laminar structure and contains a heterogeneous group of neurons that differ in gene expression, morphology, connectivity, and electrophysiological properties (Sheets et al., 2011; Oswald et al., 2013; Shepherd, 2013; Suter et al., 2013; Economo et al., 2018). These neurons can be classified into the intratelencephalic neurons (ITNs) distributed across the layers (L) 2-6 and the pyramidal tract neurons (PTNs) that mainly locate within the L5b (Oswald et al., 2013; Shepherd, 2013). Both PTNs and ITNs project to the striatum and receive basal ganglia feedbacks through the transition of the motor thalamus (Bodor et al., 2008; Kita and Kita, 2011; Kress et al., 2013; Lee et al., 2020). In physiological state, M1 network dynamics plays an essential role in the execution and coordination of complex movements and the acquisition of motor skills (Guo et al., 2015a; Kawai et al., 2015; Sreenivasan et al., 2016; Barthas and Kwan, 2017; Ebbesen et al., 2017; Wang et al., 2017; Economo et al., 2018; Sauerbrei et al., 2020). In parkinsonian state, M1 exhibits an aberrant oscillation and bursting pattern of activity at both individual neuron and population levels (Goldberg et al., 2002; Mallet et al., 2008; Pasquereau and Turner, 2011; Shimamoto et al., 2013; Hemptinne et al., 2015; Pasquereau et al., 2016), which disrupt its normal function in motor control and motor learning.

The abnormal neuronal activity in the M1 has been hypothesized to be a consequence of pathological basal ganglia output (Hosp et al., 2011, 2015; Pasquereau and Turner, 2011; Guo et al., 2015b; Pasquereau et al., 2016), but the cellular adaptations could also play a role in such abnormal activities. Moreover, compelling evidence suggests that PTNs and ITNs adapt differently to the degeneration of midbrain DA neurons (Pasquereau and Turner, 2011; Pasquereau et al., 2016).Yet, the cellular and synaptic mechanisms of such adaptations remain unknown. We hypothesized that loss of SN DA neurons alters the intrinsic properties of M1 pyramidal neurons in a cell-type-specific manner, which can contribute to the abnormal neuronal activity of M1 in parkinsonism. We addressed these hypotheses using electrophysiology and retrograde tracing approach in mice with 6-hydroxydopamine (6-OHDA) lesion, an established model of parkinsonism.

## Material and Methods

### Animals

Adult (3–4 months old) C57BL/6 mice of both sexes were used in the study. Mice were housed up to four animals per cage under a 12-h light/12-h dark cycle with access to food and water *ad libitum* in accordance with Van Andel Institute IACUC and NIH guidelines for care and use of animals. All the animal experiments were reviewed and approved by IACUC at Van Andel Institute.

### Stereotaxic surgery

Mice, under 2% isoflurane anesthesia, were placed in a stereotaxic frame (Kopf), and were supported by a thermostatic heating pad. To induce degeneration of SN DA neurons, 6-OHDA (3–4 mg/mL, 1.0 μL) was injected unilaterally into the medial forebrain bundle (MFB, from bregma (in mm): anteroposterior, −0.7, mediolateral, +1.2, dorsoventral, −4.7) over 10 min using a 10-μl syringe (Hamilton) and a motorized microinjector (Stoelting, Wood Dale, IL, USA) (Chu et al., 2017). Controls were injected in the same location with vehicle. Desipramine (25 mg/kg) and pargyline (50 mg/kg) were subcutaneously injected 30–40 min prior to 6-OHDA injection, to enhance the toxicity of 6-OHDA on dopaminergic neurons and reduce damages to noradrenergic system (Chu et al., 2015, 2017). To label and identify PTNs and ITNs in the M1, animals also received red or green Retrobeads (0.3 μL, Lumafluor Inc) injected into the ipsilateral pontine nuclei (from bregma (in mm): anteroposterior, −5.0; mediolateral, +0.6; dorsoventral, −5.0; 0.3 μl per injection) or the contralateral dorsolateral striatum (from bregma (in mm): anteroposterior, +0.4; mediolateral, −2.0; dorsoventral, −2.8). The surgical wound was closed using sutures and mice were allowed to recover after surgery in a heated cage with access to food and water at the cage-floor level. Coronal brain sections containing the forelimb region of M1 from 6-OHDA- or vehicle-injected mice were prepared 3-4 weeks after surgery for *ex vivo* electrophysiology.

### Slice preparation

Mice were deeply anesthetized with intraperitoneal avertin (250–300 mg/kg) and then were perfused transcardially with ice-cold, sucrose-based, artificial cerebrospinal fluid (aCSF) containing (in mM) 230 sucrose, 26 NaHCO_3_, 10 glucose, 10 MgSO_4_, 2.5 KCl, 1.25 NaH_2_PO_4_, and 0.5 CaCl_2_. Next, coronal brain slices (250 μm) containing M1 forelimb regions were prepared in the same slicing solution using a vibratome (VT1200S; Leica Microsystems Inc., Buffalo Grove, IL, USA). Brain slices were kept in normal aCSF (in mM: 126 NaCl, 26 NaHCO_3_, 10 glucose, 2.5 KCl, 2 CaCl_2_, 2 MgSO_4_, 1.25 NaH_2_PO_4_, 1 sodium pyruvate and 0.005 L-glutathione) equilibrated with 95% O_2_ and 5% CO_2_ for 30 min at 35 °C and then held at room temperature for 30 min. Electrophysiological recordings started after this 1-hour recovery period.

### *Ex vivo* electrophysiology recording

Brain slices were transferred into a recording chamber perfused at a rate of 4 mL/min with synthetic interstitial fluid (in mM: 126 NaCl, 26 NaHCO_3_, 10 glucose, 3 KCl, 1.6 CaCl_2_, 1.5 MgSO_4_, 1.25 NaH_2_PO_4_) equilibrated with 95% O_2_ and 5% CO_2_ at 35 °C via a feedback-controlled in-line heater (TC-324C, Warner Instruments). DNQX (20 μM), D-APV (50 μM), and SR-95531 (10 μM) were routinely added to block synaptic transmission mediated by ionotropic glutamatergic and GABAergic receptors. Neurons were visualized and recorded under gradient contrast SliceScope 6000 (Scientifica, UK) with infrared illumination using a CCD camera (SciCam Pro, Scientifica, UK) and motorized micromanipulators (Scientifica, UK). Individual neurons labeled with Retrobeads in the M1 L5b were identified using a 60X water immersion objective lens (Olympus, Japan) and targeted for whole-cell patch-clamp recording, using a MultiClamp 700B amplifier and a Digidata 1550B digitizer under the control of pClamp11 (Molecular Devices, San Jose, USA). Data were collected at a sampling rate of 20-50 KHz. Borosilicate glass pipettes (O.D. = 1.5 mm, I.D. = 0.86 mm, item#: BF150-86-10, Sutter Instruments, Novato, CA) for patch clamp recordings (4–7 MΩ) were pulled using a micropipette puller (P1000, Sutter Instruments, Novato, CA). Pipette capacitance was compensated for before the formation of the whole-cell configuration. Series resistance (Rs) was regularly monitored and compensated for with the bridge balance circuit in current clamp mode. Liquid junction potential (about 11 mV) was corrected. Under current clamp mode, pipette capacitance was neutralized using MultiClamp 700B amplifier. Briefly, ~10 mV sawtooth pattern of voltage trace was generated through “tuning” function. Then, the degree of capacitance neutralization was determined by gradually increasing capacitance values until voltage oscillations were about to onset.

Retrobeads-labeled neurons in the L5b of M1 were recorded in whole-cell current-clamp mode to study their intrinsic properties. In these experiments, glass pipettes were filled with a potassium gluconate-based internal solution of (in mM) 140 K-gluconate, 3.8 NaCl, 1 MgCl_2_, 10 HEPES, 0.1 Na_4_-EGTA, 2 ATP-Mg, and 0.1 GTP-Na, pH 7.3, osmolarity 290 mOsm. The resting membrane potential (V_m_) was recorded once the whole-cell configuration was obtained. The intrinsic properties and excitability of M1 pyramidal neuron subtypes were studied by injecting a family of current steps ranging from −240 pA to 720 pA in 40-pA increments and with a duration of 1 s. Current injections were from V_m_ and no additional holding current was injected. Input resistance (R_m_) was determined by measuring the steady-state voltage responses to a series of 1-s hyperpolarizing currents, as the slope of linear fit to the resulting voltage-current relationship. The membrane time constant (τ_m_) was determined as the slow component of a double-exponential fit of the voltage decays in response to −40 or −80 pA, 1-s current injections (i.e., a 5–10 mV voltage change). Cell capacitance (C_m_) was derived from the equation C_m_ = τ_m_ /R_m_.

The rheobase, as an indicator of neuronal excitability, was defined as the current intensity to elicit an action potential (AP). Rheobase was determined by first probing the response of neurons to 1-s current injections to define the targeted range of current intensities, and then by bounding the rheobase through small current steps (1 pA increments). AP waveforms at the rheobase were systematically analyzed and quantified using Clampfit 11.1. Specifically, the threshold of AP was determined as the voltage level at which d*V*/d*t* exceeded 20 mV/ms. AP amplitude was defined as the voltage difference between the threshold and the peak voltage. AP half-width was measured as the time difference at 50% of AP amplitude. Fast afterhyperpolarizations (fAHPs) were measured as the negative voltage peaks relative to the threshold within 2–5 ms from AP threshold (Villalobos et al., 2004; Bean, 2007). To quantify spike trains, spike frequency adaptation was measured as the ratio between the last inter-spike-interval and the average of the first two inter-spike-intervals. The gain of the spike-current curve was defined as the slope of the linear portion of the instantaneous spike-current curve.

### Immunohistochemistry

Immunoreactivity of tyrosine hydroxylase (TH) and norepinephrine transporter (NET) was assessed to validate the degeneration of the nigrostriatal DA projections and noradrenergic projections to M1 in 6-OHDA-injected mice. Briefly, brain tissues were first fixed in 4% paraformaldehyde in 0.1 M phosphate buffer, pH7.4, at 4 °C for 12 hours before rinsing in phosphate-buffered saline (PBS, 0.05 M; pH 7.4). Tissues was re-sectioned at 70 μm using a vibratome (Leica VT1000S; Microsystems Inc.). Immunochemical detection of TH was conducted in PBS containing 0.2% Triton X-100 (Fisher Scientific) and 2% normal donkey serum (Sigma-Aldrich, St. Louis, MO). Brain sections were incubated in primary antibody (mouse anti-TH antibody, 1:2000, cat#: MAB318, Sigma-Aldrich, St. Louis, MO, USA; or mouse anti-NET antibody, 1:1000, cat# 211463, Abcam, Cambridge, MA, USA) for 48 h at 4 °C or overnight at room temperature (RT), washed in PBS, and then incubated in the secondary antibody (donkey anti-mouse Alexa Fluor 488, cat#: 715-545-150 or donkey anti-mouse Alexa Fluor 594, cat#715-585-150; 1: 500, Jackson ImmunoReseach, West Grove, PA, USA) for 90 minutes at RT before washing with PBS. Brain sections were mounted on glass slides using VECTASHIELD antifade mounting medium (cat#: H-1000, Vector Laboratories, Burlingame, CA, USA) and were cover-slipped. TH immunoreactivity in the dorsal striatum and the overlying motor cortex in the same section was imaged using an Olympus BX63F microscope equipped with an Olympus DP80 camera or a confocal laser scanning microscope (A1R; Nikon, Melville, USA). TH immunoreactivity of each section was determined as the difference between the immunofluorescence between the dorsal striatum and the motor cortex (Chu et al., 2015). The intensity of TH and NET immunofluorescence was quantified using ImageJ (NIH, Bethesda, MD, USA).

### Animal Behavior

To validate the development of parkinsonian bradykinesia and akinesia prior to electrophysiology studies, vehicle- and 6-OHDA-injected mice were subject to 1) open field locomotion test for 10 min and their locomotor activity as well as rotations were monitored and quantified using Anymaze software (Stoelting, Wood Dale, IL); 2) cylinder test to assess their spontaneous forelimbs use during weight-bearing touch on wall of a 600 ml glass cylinder. Spontaneous exploration of mice was recorded using an HD digital camcorder at 60 fps (Panasonic, HC-V180K) and analyzed off-line by a researcher blinded to treatments.

### Experimental design and statistical analysis

Data were analyzed in Clampfit 11.1 (Molecular Devices, San Jose, CA) and ImageJ (NIH). Statistics were done using Prism8 (GraphPad Software, San Diego, CA, USA). To minimize the assumption of the data normality, we used non-parametric, distribution-independent Mann-Whiney U (MWU) or Wilcoxon signed rank (WSR) tests for non-paired or paired data comparisons, respectively. The Holm-Bonferroni correction was applied for multiple comparisons. All tests were two-tailed, and an α-level of 0.05 was used to determine statistical significance. Results are reported as median and interquartile range. Boxplots illustrate the median (central line), interquartile range (box), and 10–90% range (whiskers) of data. Data are available upon request.

## Results

All physiology studies were conducted between 3- and 4-weeks post-surgery, when the level of midbrain DA degeneration and functional adaptations in the brain have reached maximum and been stabilized (Vila et al., 2000; Viaro et al., 2011). At 3 weeks post-surgery, mice with 6-OHDA lesion (hereinafter, “6-OHDA mice”) showed featured parkinsonians akinesia and bradykinesia versus vehicle-injected controls (hereinafter, “controls”), as supported by 1) a reduced travelled distance in open-field locomotion test (6-OHDA mice = 18 [16, 22] m, controls = 41 [32, 50] m, n = 13 mice from each group, p < 0.0001, MWU); 2) ipsilateral rotations (%ipsilateral rotations, 6-OHDA mice = 100 [97, 100]%, controls = 41 [26, 53]%, n = 13 mice from each group, p < 0.0001, MWU); and impaired forelimb use (%ipsilateral forelimb use: 6-OHDA mice = 88 [83, 92]%, controls = 50 [49, 53]%, n = 13 mice from each group, p < 0.0001, MWU). In *post hoc* histology studies, 6-OHDA mice showed > 80% reduction of striatal TH immunoreactivity (TH-ir) in the hemisphere with lesion (%TH-ir in the ipsilateral versus the contralateral striatum, 6-OHDA mice = 0 [0, 0]%, and controls = 97 [85, 102]%, n = 16 mice for each group). In addition, desipramine was routinely administered to inhibit reuptake of 6-OHDA by noradrenergic terminals. Given the heavy noradrenergic innervation of the cortical regions, we therefore validated the effect of desipramine by immunostaining of cortical NET and striatal TH in a separate cohort of animals. %NET-ir in the ipsi- versus contra-lateral L5 of M1, 6-OHDA = 92 [78, 108]%, while %TH-ir in the ipsi- versus contra-lateral striatum = 0 [0, 3.5]%, n = 8 sections/4 mice.

### Cell-type-specific decrease in the intrinsic excitability of M1 L5b pyramidal neurons following the loss of midbrain DA neurons

We compared the intrinsic excitability of projection-defined pyramidal neurons in the L5b of M1 from controls and 6-OHDA mice. To identify M1 pyramidal neurons based on long-range projections, Retrobeads were stereotaxically injected into the ipsilateral pontine nuclei (Figure 1A) and the contralateral striatum (Figure 2A) to retrogradely label PTNs (Figure 1B) and ITNs (Figure 2B), respectively. The intrinsic electrophysiological properties of M1 PTNs and ITNs were assessed using whole-cell current-clamp recording in the presence of antagonists of ionotropic glutamatergic and GABAergic receptors, including DNQX (20 μM), D-APV (50 μM), and SR95531 (10 μM).

**Figure 1.**
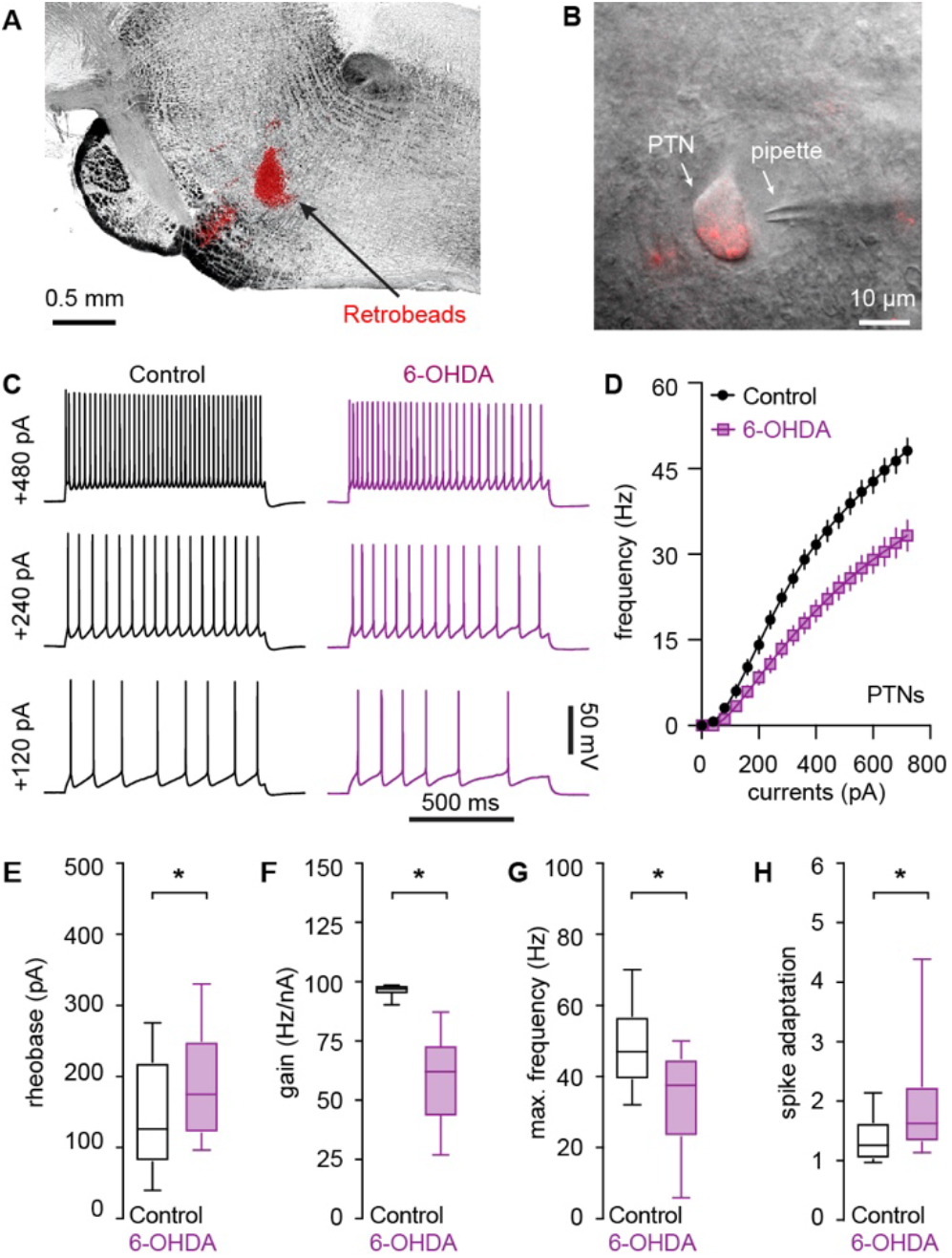
Decreased intrinsic excitability of PTNs in the L5b of M1 following the loss of midbrain DA neurons. **A and B**) Representative graphs showing the injection site of red Retrobeads in the pontine nuclei (**A**) and a retrogradely labeled PTN in L5b of M1 (**B**). **C and D**) Representative spike trains of PTNs evoked by somatic current injections from controls and 6-OHDA mice (**C**), and the frequency-current relationship of the PTNs from controls and 6-OHDA mice (**D**). **E-H**) Summarized results showing an increased rheobase (**E**), a reduced gain of spike-current curve (**F**), a decreased firing frequency in response to maximal current injections (i.e., 720 pA) (**G**), and an enhanced spike adaptation (**H**) in PTNs from 6-OHDA mice relative to those from controls. *, p < 0.05, MWU.

**Figure 2.**
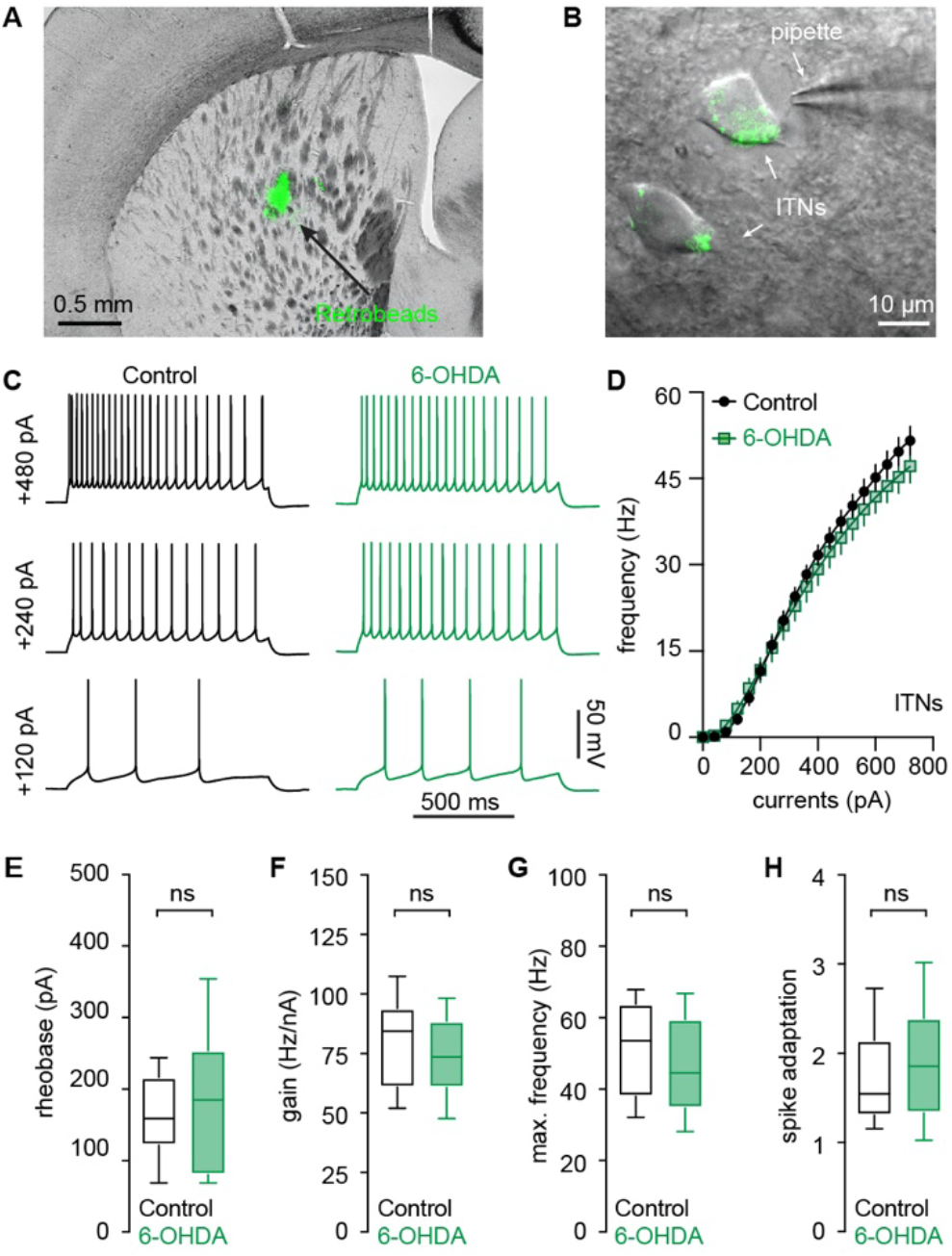
Intact intrinsic excitability of ITNs in the M1 L5b following the loss of midbrain DA neurons. **A and B**) Representative graphs showing the injection site of green Retrobeads in the dorsal striatum (**A**) and retrogradely labeled M1 L5b ITNs (**B**). **C and D**) Representative spikes trains evoked by somatic current injections into ITNs from control and 6-OHDA mice (**C**), and the frequency-current relationship of ITNs from controls and 6-OHDA mice (**D**). **E-H**) Boxplots showing a similar rheobase (**E**), an unaltered gain of spike-current curve (**F**), firing frequency in response to maximal current injections (i.e., 720 pA) (**G**), and spike adaptation (**H**) between ITNs from 6-OHDA mice and controls. ns, not significant, MWU.

The intrinsic excitability of M1 pyramidal neuron subtypes was assessed by somatic injection of a range of depolarizing currents from V_m_ in controls and 6-OHDA mice (Figures 1C, D). First, the intensity of somatic current injection to elicit single action potential (AP) increased significantly for PTNs from 6-OHDA mice relative to controls (rheobase, control = 126 [80, 220] pA), n = 39 neurons/4 mice, 6-OHDA = 175 [120, 250] pA, n = 30 neurons/3 mice, p = 0.03, MWU, Figure 1E). Moreover, the gain of spike-current curve decreased significantly in M1 PTNs from 6-OHDA mice relative to controls (control = 97 [95, 98] Hz/nA, n = 39 neurons/4 mice; 6-OHDA = 62 [44, 73] Hz/nA, n = 30 neurons/3 mice; p < 0.0001, MWU; Figure 1F). The same reduction was also detected for the maximal spike frequency (e.g., frequency at 720 pA current injection, control = 47 [39, 57] Hz, n = 39 neurons/4 mice; 6-OHDA = 38 [23, 45] Hz, n = 30 neurons/3 mice; p = 0.0002, MWU; Figure 1G). M1 PTNs could sustain persistent firing with minimal spike adaptation during repetitive firing in the physiological state (Figure 1C) (Oswald et al., 2013; Suter et al., 2013). However, PTNs showed significantly enhanced spike adaptation in 6-OHDA mice (Figure 1 C). The spike adaptation of a 10-Hz spike train, control = 1.2 [1.0, 1.6], n = 39 neurons/4 mice; 6-OHDA = 1.6 [1.3, 2.3], n = 30 neurons/3 mice; p = 0.0011, MWU (Figures 1C, H). Altogether, these results suggest a significant reduction in the intrinsic excitability of M1 PTNs in parkinsonism.

In contrast, M1 ITNs from controls and 6-OHDA mice showed comparable intrinsic excitability (Figure 2). This was reflected by four measures, namely, a) similar rheobase (control = 155 [118, 213] pA, 6-OHDA = 184 [79, 252] pA, n = 30 neurons/3 mice for each group, p = 0.4, MWU, Figure 2E); b) the unaltered gain of spike-current curve (control = 84 [61, 94] Hz/nA, 6-OHDA = 74 [61, 89] Hz/nA, n= 30 neurons/3 mice for each group; p = 0.3, MWU; Figure 2F); c) the maximal spike frequency (control = 53 [38, 64] Hz, 6-OHDA = 45 [35, 60] Hz, n = 30 neurons/3 mice for each group; p = 0.15, MWU; Figure 2D, G); and d) spike adaptations (control = 1.6 [1.3, 2.2], 6-OHDA = 1.9 [1.3, 2.4], n = 30 neurons/3 mice for each group; p = 0.39, MWU; Figure 2C, H). Altogether, our results thus suggest that the M1 L5b pyramidal neurons exhibited cell-type-specific decrease of intrinsic excitability in parkinsonism.

### No change in the passive membrane properties of M1 L5b pyramidal neurons following the loss of midbrain DA neurons

Next, we determined whether alterations in the passive membrane properties contribute to the difference in the intrinsic excitability of M1 PTNs and ITNs in parkinsonism. In the presence of antagonists of fast ionotropic glutamatergic and GABAergic receptors, PTNs in 6-OHDA mice and controls had similar resting membrane potential (V_m_, control = −75.8 [−77.2, −74.9] mV, n = 39 neurons/4 mice; 6-OHDA = −76.1 [−77.1, −74.9] mV, n= 30 neurons/3 mice; p = 0.87, MWU, Figure 3A) and input resistance (R_m_, control = 72.9 [60, 83.8] MΩ, n = 39 neurons/4 mice; 6-OHDA = 71.3 [64, 86.6] MΩ, n= 30 neurons/3 mice; p = 0.91, MWU, Figure 3B). In addition, PTNs from 6-OHDA mice did not show changes in their cell capacitance (C_m_) and membrane time constant (τ_m_) relative to those from controls (C_m_, control = 212 [160, 239] pF, n = 39 neurons/4 mice, 6-OHDA = 192 [167, 268] pF, n = 30 neurons/3 mice, p = 0.7, MWU, Figure 3C; τ_m_, control = 19.4 [14.4, 23.4] ms, n = 39 neurons/4 mice, 6-OHDA = 20.9 [12.4, 30.5] ms, n = 30 neurons/3 mice, p = 0.5, MWU, Figure 3D).

**Figure 3.**
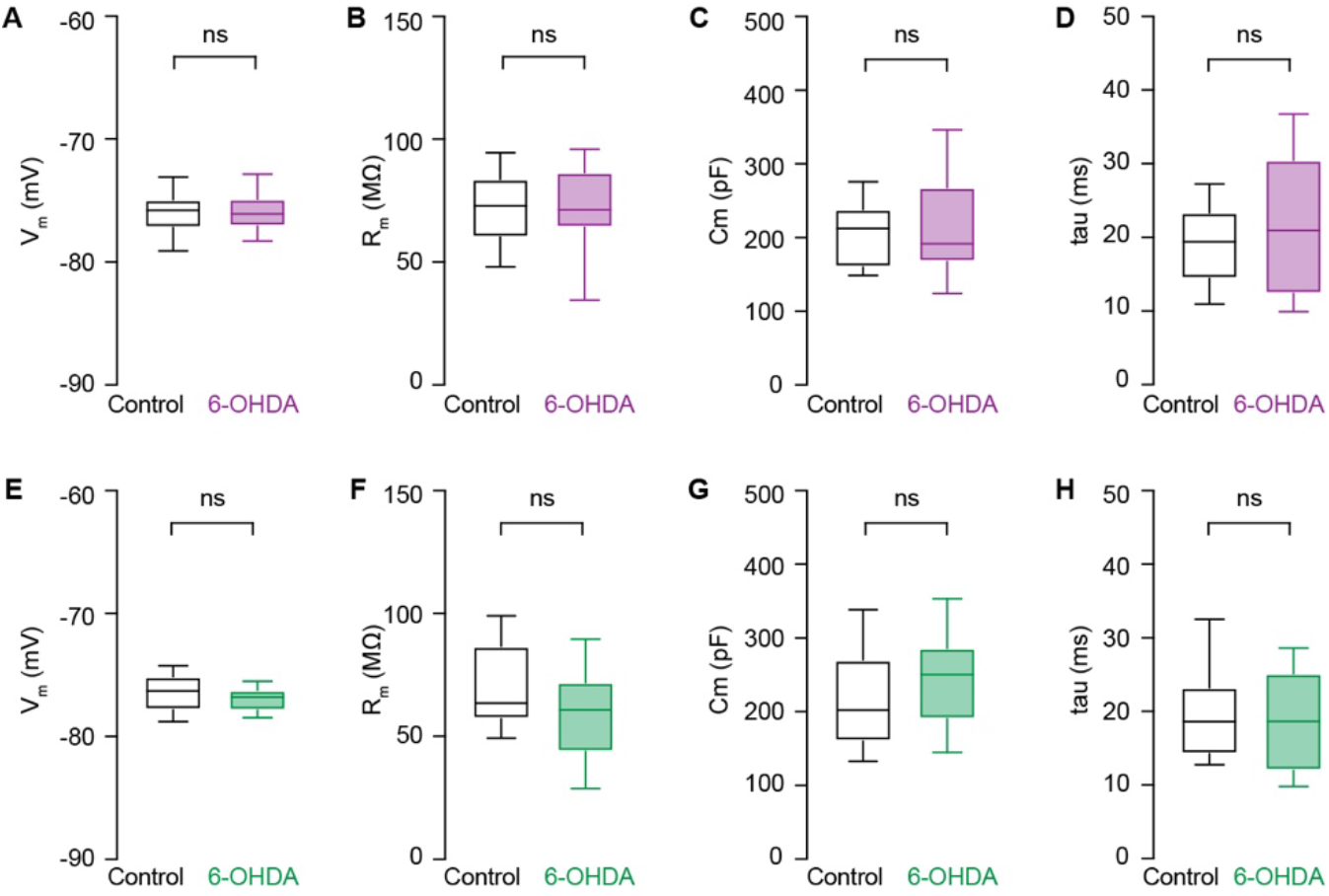
No change in the passive membrane properties of PTNs and ITNs in the M1 L5b following the loss of midbrain DA neurons. **A-D**) Boxplots showing no changes in the resting membrane potential (**A**), input resistance (**B**), cell capacitance (**C**), and time constant (**D**) of PTNs from 6-OHDA mice relative to those from controls. **E-H**) Similar measures of passive membrane properties of ITNs from controls and 6-OHDA mice. ns, not significant, MWU.

Similarly, ITNs showed comparable passive membrane properties between groups, namely, V_m_ (control = −76.3 [−77.8, −75.1] mV, 6-OHDA = −76.8 [−77.9, −76.2] mV, n = 30 neurons/3 mice for each group, p = 0.2, MWU, Figure 3E), R_m_ (control = 63.5 [57.1, 86.7] MΩ, 6-OHDA = 60.8 [43.8, 72] MΩ, n = 30 neurons/3 mice for each group, p = 0.15, MWU, Figure 3F), C_m_ (control = 202 [160, 270] pF, 6-OHDA = 250 [190, 287] pF, n = 30 neurons/3 mice for each group, p = 0.13, MWU, Figure 3G), and τ_m_ (control = 18.6 [14.2, 23.3] ms, 6-OHDA = 18.6 [12, 25.2] ms, n = 30 neurons/3 mice for each group, p = 0.6, MWU, Figure 3H). Altogether, these results suggest that the cell-type-specific decrease in the excitability of M1 pyramidal neurons is not due to alterations in their passive membrane properties.

### Cell-type-specific changes in action potential waveforms of M1 L5b pyramidal neurons following the loss of midbrain DA neurons

The shape of the action potential is determined by the interaction of numerous voltage-gated ion channels and has profound effects on the rate and pattern of neuronal firing (Bean, 2007). We therefore analyzed the morphology of AP of PTNs and ITNs at the rheobase from controls and 6-OHDA mice (Figures 4A, H). In parkinsonian animals, M1 PTNs showed a depolarized AP threshold (controls = −49.7 [−51.5, −47.7] mV, n = 39 neurons/4 mice, 6-OHDA = −47.9 [−50.3, −45.8] mV, n = 30 neurons/3 mice, p = 0.015, MWU, Figure 4B) and a striking AP broadening (AP half-width, controls = 0.69 [0.66, 0.75] ms, n = 39 neurons/4 mice; 6-OHDA = 0.77 [0.72, 0.82] ms, n = 30 neurons/3 mice; p = 0.0008, MWU, Figure 4C). The broadened AP of PTNs was associated with a significantly prolonged AP rise time (controls = 0.19 [0.18, 0.22] ms, n = 39 neurons/4 mice; 6-OHDA = 0.21 [0.2, 0.24] ms, n = 30 neurons/3 mice; p = 0.0024, MWU; Figures 4D) and also a prolonged decay time (controls = 0.8 [0.69, 0.84] ms, n = 39 neurons/4 mice, 6-OHDA = 0.9 [0.76, 0.97] ms, 30 neurons/3 mice; p = 0.0023, MWU; Figure 4E). In parkinsonian animals, M1 PTNs also exhibited significant decreases in the maximal rates of AP depolarization (the rise rate, controls = 434 [398, 493] mV/ms, n = 39 neurons/4 mice, 6-OHDA = 390 [340, 443] mV/ms, n= 30 neurons/3 mice, p = 0.004, MWU, Figure 4F) and repolarization (the decay rate, controls = −102 [−87, −111] mV/ms, n = 39 neurons/4 mice, and 6-OHDA = −90 [−81, −99] mV/ms, n = 30 neurons/3 mice, p = 0.012, MWU, Figure 4G). Thus, the decreased excitability of PTNs in parkinsonism involves impairments in both Na^+^ and K^+^ conductance that underlies AP depolarization and repolarization, respectively (Bean, 2007).

**Figure 4.**
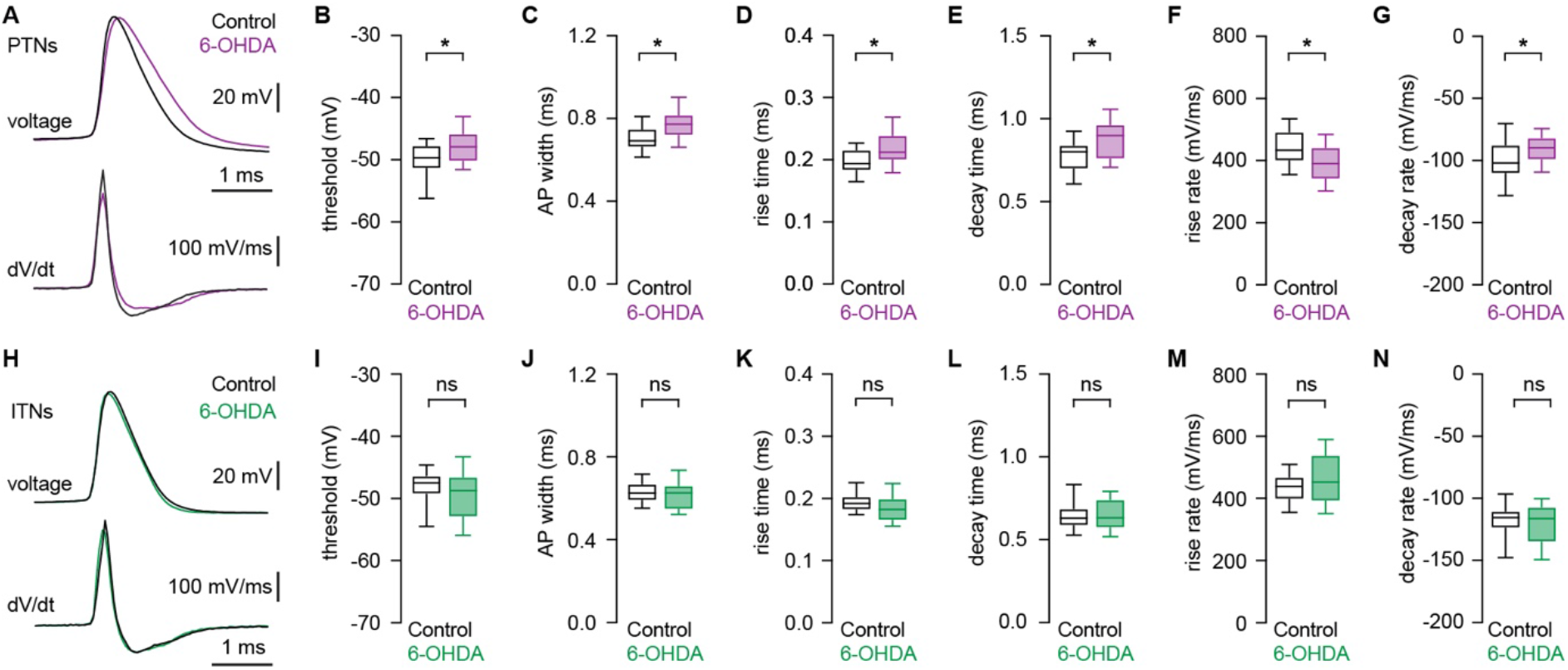
Distinct changes in AP waveforms of PTNs and ITNs in the M1 L5b following loss of midbrain DA neurons. **A**) **Top**, representative AP waveforms of PTNs from controls and 6-OHDA mice. APs were aligned at threshold and overlaid for comparison. **Bottom**, the corresponding *d*V/*d*t traces plotted against time, showing depolarizing and repolarizing rates of APs. **B and C**) Boxplots showing AP threshold and width of PTNs in 6-OHDA mice and controls. **D and E**), Boxplots showing prolonged AP rise time (**D**) and decay time (**E**) in PTNs from 6-OHDA mice relative to those from controls. **F and G**), Boxplots showing decreased AP rise rate (**F**) and decay rate (**G**) in PTNs from 6-OHDA mice relative to those from controls. **H**) **Top**, representative AP waveforms of ITNs from controls and 6-OHDA mice. APs were aligned at threshold and overlaid for comparison. **Bottom**, the corresponding *d*V/*d*t traces plotted against time, showing depolarizing and repolarizing rates of APs. **I-N**) Boxplot showing similar AP threshold (I) and width (J), rise (K) and decay (L) time, as well as rise (M) and decay (N) rates in ITNs from 6-OHDA mice relative to those from controls. ns, not significant; *, p < 0.05, MWU.

In contrast, ITNs from controls and 6-OHDA mice showed unaltered AP threshold (controls = −47.5 [−49.3, −46.3] mV, 6-OHDA = −48.7 [−53, −46.5] mV, n = 30 neurons/3 mice for each group, p = 0.24, MWU, Figure 4I) and AP width (controls = 0.63 [0.59, 0.67] ms; 6-OHDA = 0.63 [0.55, 0.66] ms, n = 30 neurons/3 mice for each group; p = 0.56, MWU; Figure 4J). The similar AP width of ITNs was associated with unaltered time and rates of AP depolarization and repolarization. The rise time was controls = 0.19 [0.18, 0.2] ms, and 6-OHDA = 0.18 [0.16, 0.2] ms (p = 0.13, MWU, Figure 4K). The decay time was controls = 0.63 [0.58, 0.69] ms, and 6-OHDA = 0.63 [0.57, .074] ms (p = 0.94, MWU, Figure 4L). The rise rate values were, controls = 438 [396, 469] mV/ms, and 6-OHDA = 452 [390, 541] mV/ms (p = 0.3, MWU, Figure 4M), and the decay rate values were, controls = −115 [−110, −124] mV/ms, and 6-OHDA = −116 [−107, −135] mV/ms (p = 0.67, MWU, Figure 4N).

Altogether, these results suggest that disrupted ionic mechanisms for AP generation contribute to cell-type-specific decrease in the intrinsic excitability of PTNs in parkinsonism.

### Progressive alterations in AP waveforms of PTNs during repetitive firing following the loss of SN DA neurons

The literature suggests that M1 PTNs are capable to sustain constant AP waveform during repetitive firing (Suter et al., 2013). Thus, we elicited spike trains containing 10 APs from M1 PTNs in controls and 6-OHDA mice and analyzed waveforms of the first and the last APs to study activity-dependent AP modulation during repetitive firing. We noticed a stronger activity-dependent AP waveform modulation in PTNs from 6-OHDA mice relative to those from controls (Figure 5A, B). Specifically, PTNs from both groups exhibited an increased AP threshold during repetitive firing (threshold in controls, 1st AP= −51.8 [−54.8, −50.5] mV, last AP = −50.7 [−52.7, −47.8] mV, n = 25 neurons/5 mice, p < 0.001, WSR; threshold in 6-OHDA mice, 1st AP = −51 [−52.5, −47.1] mV, last AP = −47.2 [−48.8, −45.6] mV, n= 25 neurons/5 mice, p < 0.001, WSR; Figure 5C). Repetitive firing leads to a significantly depolarized threshold of the last APs from 6-OHDA mice relative to those from controls (p = 0.0003 between groups, MWU, Figure 5C), in addition to the elevated threshold of the 1^st^ APs (p = 0.02 between groups, MWU; Figure 5C). This is consistent with activity-dependent inactivation of voltage-dependent transient Na^+^ channels (Na_T_) during sustained firing (Bean, 2007).

**Figure 5.**
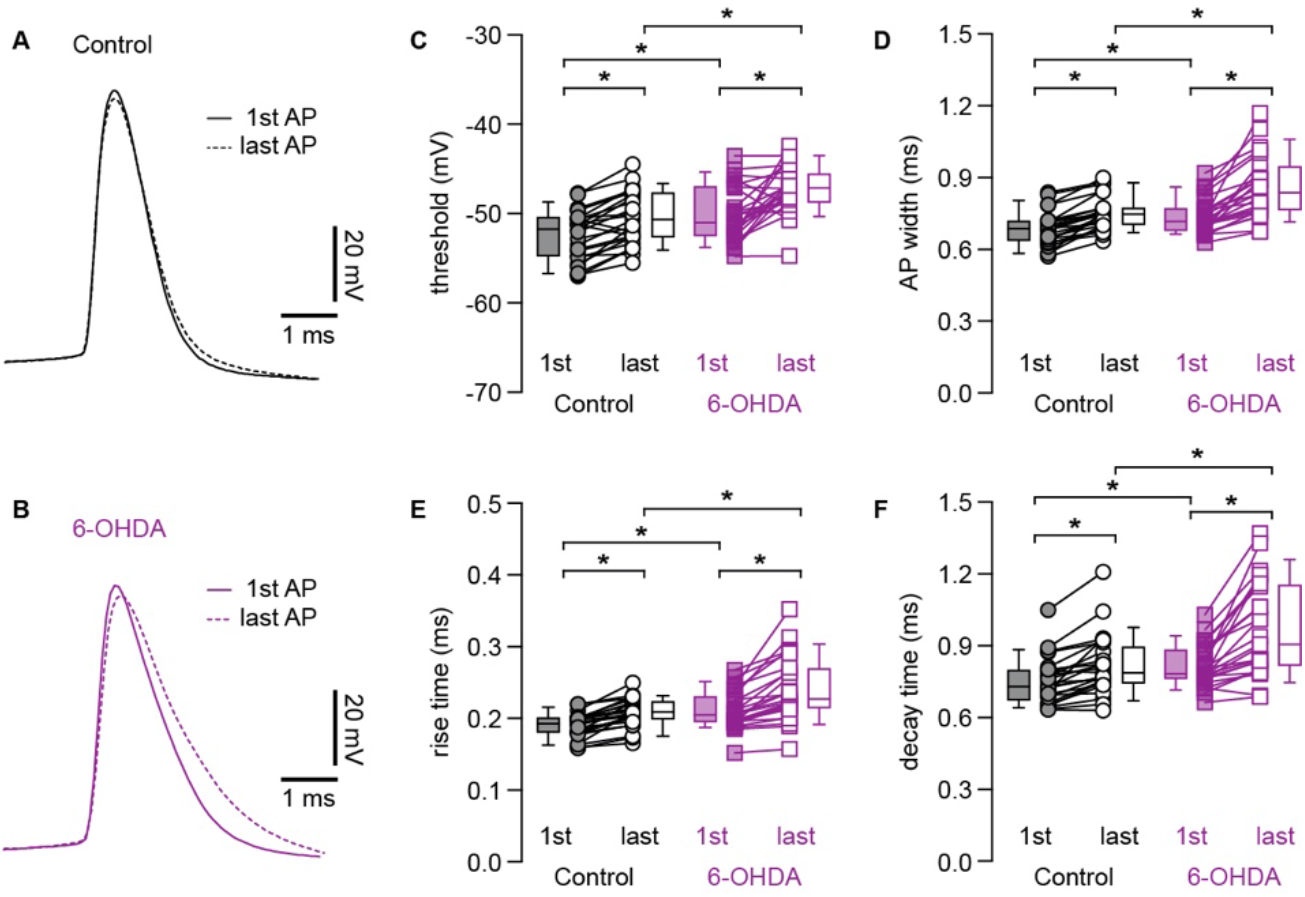
Progressive changes in AP waveforms of M1 PTNs during repetitive firing. **A-B**) The first and the 10^th^ APs from a 10-Hz spike train were superimposed for PTNs from control (**A**) or 6-OHDA mice (**B**). APs were aligned at threshold, and the solid and dashed lines indicated the first and the 10^th^ APs from the spike train, respectively. **C**) Summarized graphs showing the alterations of AP threshold between the first and the 10^th^ APs. Each line connects the AP width of the first and the 10^th^ APs from the same spike train. Box plots indicate the median plus interquartile range of the first and the 10^th^ AP. (**D-F**) Summarized graphs showing changes in AP width (D), rise time (E), and decay time (F) between the first and the 10^th^ APs in PTNs from 6-OHDA mice relative to those from controls. *, p < 0.05, WSR or MWU tests.

APs of PTNs in both controls and 6-OHDA mice also progressively broadened during repetitive firing (AP width in controls, 1^st^ AP = 0.68 [0.64, 0.72] ms, last AP = 0.75 [0.7, 0.77] ms, n= 25 neurons/5 mice, p < 0.0001, WSR; AP width in 6-OHDA mice, 1^st^ AP = 0.72 [0.68, 0.77] ms, last AP = 0.84 [0.76, 0.94] ms, n= 25 neurons/5 mice, p < 0.0001, WSR, Figure 5D). Importantly, repetitive firing led to a significantly broadened last APs in 6-OHDA mice relative to controls (width of the last AP, p = 0.0004 between groups, MWU, Figure 5D), in addition to the broadened 1^st^ AP (width of the 1^st^ AP, p = 0.01 between groups, MWU, Figure 5D). Moreover, the observed activity-dependent modulation of AP waveforms associated with prolonged rise and decay time in controls and 6-OHDA mice (Figure 5E, F). The values of AP rise time in controls were, the 1^st^ AP = 0.19 [0.18, 0.2] ms, the last AP = 0.21 [0.2, 0.22] ms (n = 25 neurons/5 mice, p < 0.0001, WSR, Figure 5E) and the values for AP rise time in 6-OHDA mice were, the 1^st^ AP = 0.2 [0.19, 0.23] ms, the last AP = 0.23 [0.21, 0.27] ms (n = 25 neurons/5 mice, p < 0.0001, WSR, Figure 5E). The rise time of the 1^st^ APs (p = 0.005 between groups, MWU, Figure 5F) and last APs (p = 0.004 between groups, MWU, Figure 5F) were both significantly increased in 6-OHDA mice relative to those in controls. Similarly, the values of AP decay time in controls were, the first AP = 0.73 [0.67, 0.8] ms, the last AP = 0.79 [0.74, 0.89] ms (n = 25 neurons/5 mice, p < 0.0001, WSR, Figure 5F). And the values AP decay time in 6-OHDA were, the first AP = 0.78 [0.76, 0.88] ms, the last AP = 0.91 [0.82, 1.15] ms (n = 25 neurons/5 mice, p < 0.0001, WSR, Figure 5F). The decay time of the 1^st^ APs (p = 0.01 between groups, MWU, Figure 5F) and the last APs (p =0.002, MWU, Figure 5F) were both significantly increased in 6-OHDA mice relative to those in controls.

Altogether, these results suggest that PTNs in parkinsonism show a stronger activity-dependent AP modulation, which, at least partially, leads to a decreased intrinsic excitability.

### Impaired function of persistent Na^+^ currents contributes to the increased threshold and decreased intrinsic excitability of PTNs in parkinsonism

Persistent sodium current (*I*_Nap_) plays an important role in determining AP threshold and its dysfunction contributes to disrupted neuronal activity in several neurological diseases (Bean, 2007; Milescu et al., 2010; Deng and Klyachko, 2016). Given the altered AP threshold and rising phase of APs in 6-OHDA mice (Figure 4B, D, F), we next examined whether *I*_Nap_ contributes to these alterations. Blockade of *I*_Nap_ by bath application of low concentration of tetrodotoxin (TTX, 20 nM) (Gage 1998; Deng 2016) significantly elevated the AP threshold at rheobase of PTNs from controls and 6-OHDA mice (controls, baseline = −50.8 [−53.7, −50] mV, TTX = −41.1 [−42.3, −38.2] mV, n = 11 neurons/3 mice, p = 0.001, WSR; 6-OHDA, baseline = −48.8 [−49.4, −46.5] mV, TTX = −41.2 [−43.5, −39.4] mV; n = 13 neurons/4 mice, p = 0.0002, WSR, Figure 6A-C). TTX (20 nM) also abolished the difference in AP threshold between groups (AP threshold, prior to TTX, p = 0.0018; in TTX, p = 0.5, MWU, Figure 6C). In contrast, blockade of *I*_Nap_ did not affect AP width at rheobase of PTNs either in controls (baseline = 0.68 [0.61, 0.72] ms, TTX = 0.7 [0.62, 0.82] ms, p = 0.2, WSR, Figure 6D) or in 6-OHDA mice (baseline = 0.8 [0.7, 0.81] ms, TTX = 0.77 [0.7, 0.83] ms, p = 0.9, WSR, Figure 6D). With the blockade of *I*_Nap_, PTNs from controls and 6-OHDA mice also exhibited activity-dependent increase in AP threshold during repetitive firing of 10 APs (controls in TTX, 1^st^ AP = −47.1 [−49.5, −40.7] mV, last AP = −40.9 [−45.8, −38.6] mV, p = 0.004, WSR, n= 9 neurons/3 mice; 6-OHDA in TTX, 1^st^ AP = −46.3 [−51.7, −43.1] mV, last AP = −41.4 [−44.8, −35.8] mV, p = 0.0005, WSR, n = 13 neurons/4 mice. Figure 6E). Further, TTX also abolished the difference in AP threshold of both the first and the last APs during repetitive firing (threshold of the first AP, p = 0.65 between groups; threshold of the last AP, p = 0.7 between groups, MWU, Figure 6E). These results indicate disrupted *I*_Nap_ function underlies the elevated AP threshold in 6-OHDA mice.

**Figure 6.**
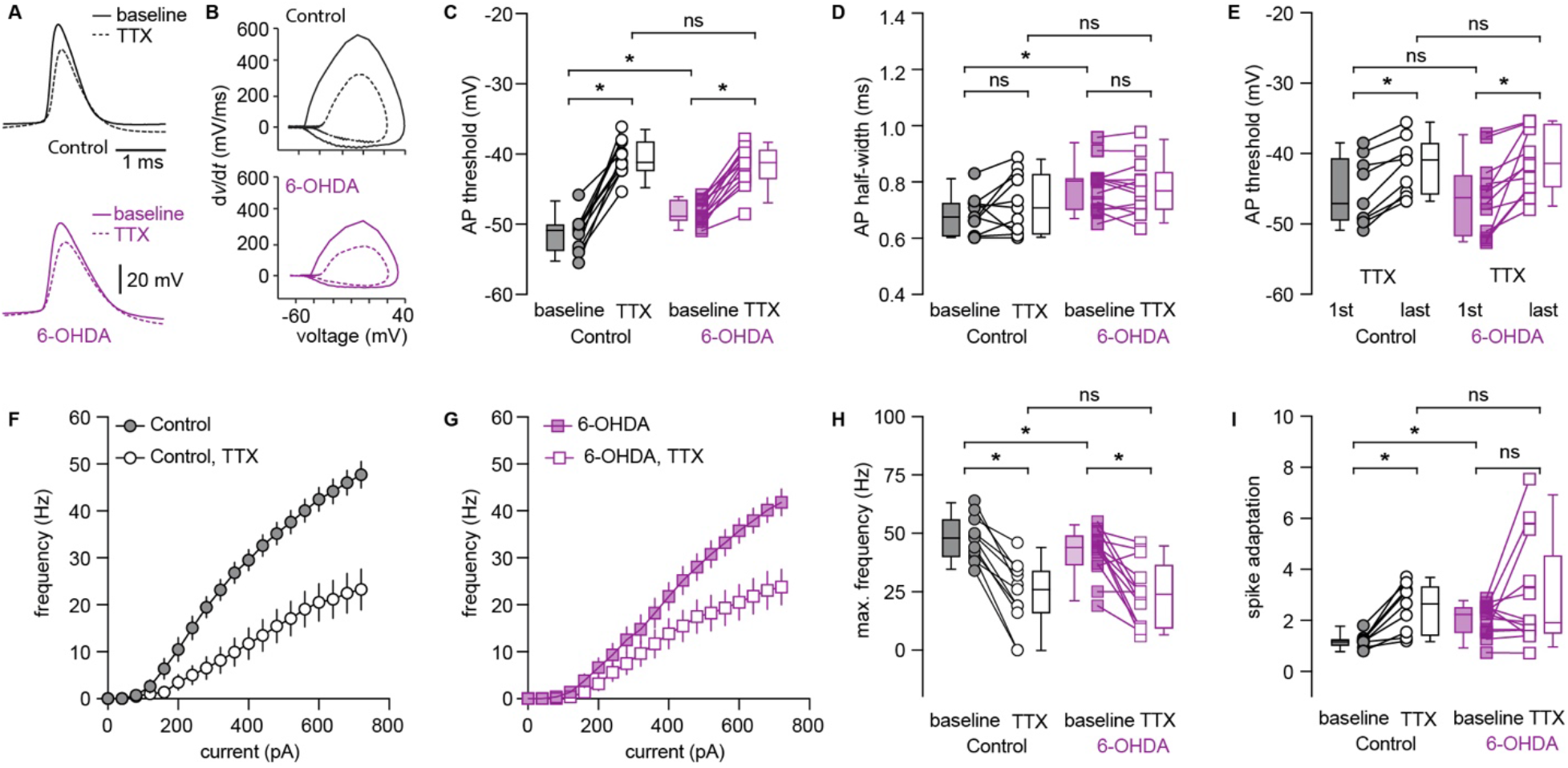
Low concentration TTX excludes the decreased intrinsic excitability of PTNs in 6-OHDA mice versus controls. **A-B**) Representative AP waveforms of PTNs at rheobase from controls (**A**, up) and 6-OHDA mice (**A**, bottom) and the corresponding phase plane plots at baseline and in the presence of TTX (20 nM). **C-D**) Summarized graphs showing changes in AP threshold (**C**) and half-width (**D**) at rheobase in the absence and presence of TTX (20 nM). **E**) Summarized graph showing threshold changes of PTNs from controls and 6-OHDA mice during repetitive firing of 10 APs in the presence of TTX (20 nM). **F-G**) Current-frequency curves of PTNs from controls and 6-OHDA mice in the absence and presence of TTX (20 nM). **H-I**) summarized graphs showing changes in the maximal firing frequency (H) and spike adaptation (**I**) of PTNs from controls and 6-OHDA mice in the absence and presence of TTX (20 nM). *, p < 0.05, n.s., not significant. WSR or MWU tests.

Next, we tested the hypothesis that a decreased basal *I*_Nap_ activity exacerbates activity-dependent Na_T_ inactivation and leads to a decreased excitability of PTNs in 6-OHDA mice. As expected, blockade of *I*_Nap_ by TTX (20 nM) significantly decreased the number of APs of PTNs from controls and 6-OHDA mice (Figure 6F, G). For example, the maximal number of APs in controls were, baseline = 48 [40, 56], TTX = 26 [16, 34] (n = 11 neurons/3 mice, p = 0.001, WSR, Figure 6H) and in 6-OHDA mice were, baseline = 44 [37, 49], TTX = 24 [10, 37] (n = 13 neurons/4 mice, p = 0.002, WSR, Figure 6H). Importantly, TTX abolished the difference in intrinsic excitability of PTN between groups (AP numbers in TTX between groups, p = 0.96, MWU, Figure 6H).

It has been reported that Na_T_ inactivation contribute to spike adaptation during repetitive firing (Fleidervish et al., 1996; Milescu et al., 2010). Consistently, blockade of *I*_Nap_ by TTX (20 nM), which promotes Na_T_ inactivation, dramatically increased spike adaptation in PTNs from controls (adaptation index, baseline = 1.18 [1.0, 1.2], TTX = 2.67 [1.43, 3.34], n = 9 neurons/3 mice, p = 0.0039, WSR, Figure 6I), but had little effect to those from 6-OHDA mice (adaptation index, baseline = 2.26 [1.55, 2.53], TTX = 1.93 [1.51, 4.55], n = 13 neurons/4 mice, p = 0.34, WSR, Figure 6I). Furthermore, TTX abolished the difference in spike adaptation between groups (controls versus 6-OHDA mice at baseline, p = 0.002; in TTX, p = 1, MWU, Figure 6I).

Thus, these data indicated that *I*_Nap_ dysfunction reduces the availability of Na_T_ for AP generation and exacerbates their inactivation during repetitive firing, leading to a decreased intrinsic excitability of PTNs in parkinsonism.

### Impaired activity of BK channels underlies the broadened APs of M1 PTNs following the loss of midbrain DA neurons

We further examined the ionic mechanisms underlying broadened APs of PTNs in 6-OHDA mice. Following the loss of midbrain DA neurons, the amplitude of the fast afterhyperpolarization (fAHP) in PTNs decreased significantly (controls = 12.5 [10.5, 14.3] mV, n = 39 neurons/4 mice; 6-OHDA = 11.2 [10.5, 11.9] mV, n = 30 neurons/3 mice; p = 0.036, MWU). These observations suggest that an altered BK channel activity probably underlies the broadened AP of PTNs in 6-OHDA mice, considering its role in AP repolarization, fAHPs, and AP broadening during neuronal repetitive firing in other brain regions (Faber and Sah, 2003; Bean, 2007; Contet et al., 2016).

Consistently, the difference in AP width at rheobase of PTNs between groups was abolished by intracellular BAPTA (10 mM, a high-affinity Ca^2+^ chelator) (controls = 0.84 [0.79, 0.92] ms, n = 26 neurons/3 mice; 6-OHDA = 0.86 [0.78, 0.94] ms, n = 26 neurons/3 mice, p = 0.84, MWU, Figure 7A, B). Moreover, in the presence of intracellular BAPTA, PTNs neurons still show activity-dependent AP broadening in both controls (1^st^ AP = 0.78 [0.68, 0.86] ms, last AP = 1.14 [0.97, 1.24] ms, n = 17 neurons/3 mice, p < 0.0001, WSR, Figure 7C) and 6-OHDA mice (1^st^ AP = 0.78 [0.73, 0.87] ms, last AP = 1.14 [0.94, 1.33] ms, n = 16 neurons/3 mice, p < 0.0001, WSR, Figure 7C). However, the extent of AP broadening during repetitive firing (10 spikes) was comparable between groups (controls versus 6-OHDA mice, half-width of the 1st AP, p = 0.5 between groups; half-width of the last AP, p = 0.8 between groups, MWU). Further, the PTNs from both groups also showed similar intrinsic excitability in the presence of BAPTA (Figure 7D). For example, the number of spikes at the maximal current injection for controls = 38 [21, 55], while the value for 6-OHDA = 29 [18, 40], n = 26 neurons /3 mice for each group; p = 0.29, MWU test.

**Figure 7.**
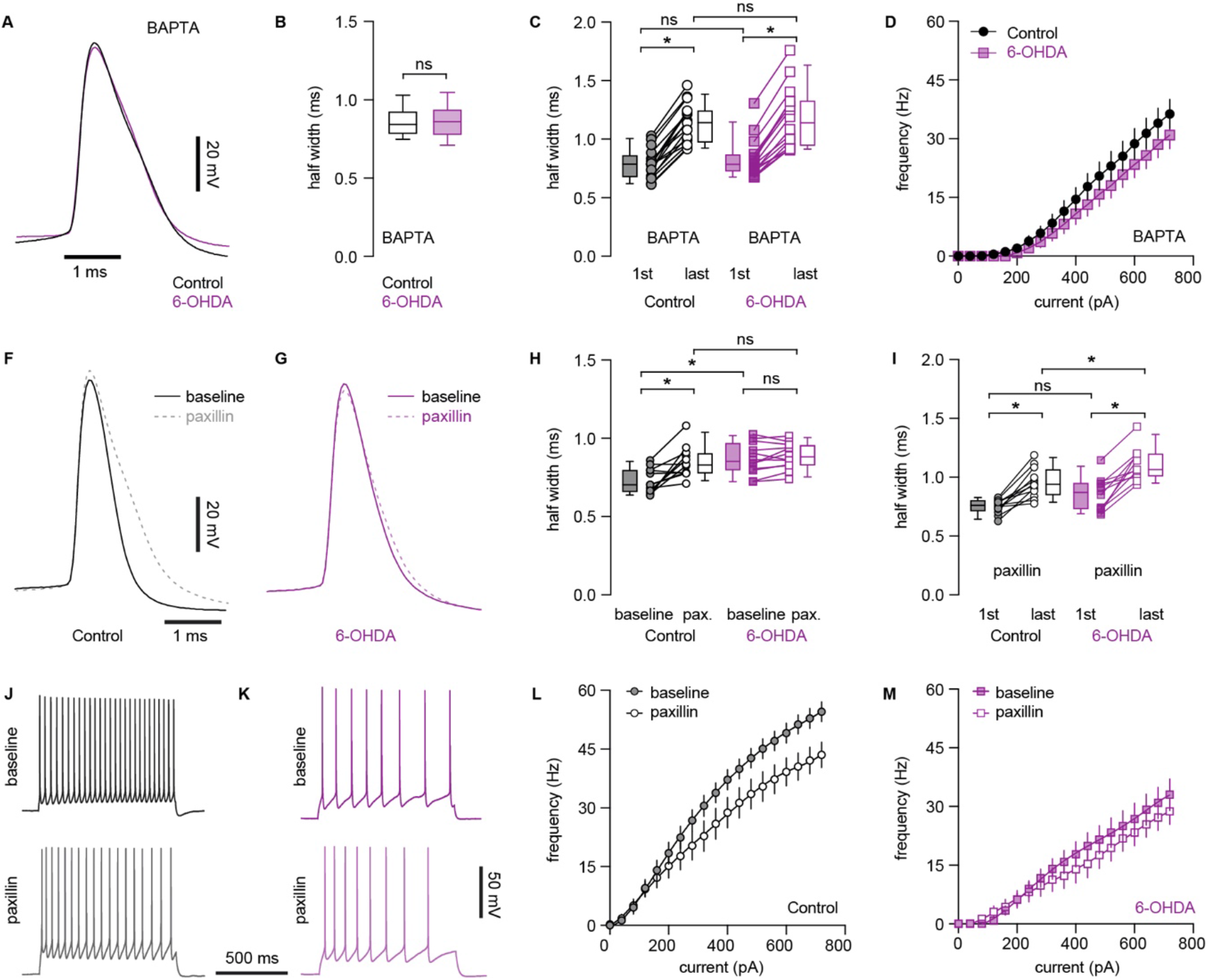
Decreased BK channel activity underlies AP broadening of PTNs following loss of midbrain DA neurons. **A**) Representative AP waveforms of PTNs from controls and 6-OHDA mice in the presence of BAPTA. APs were aligned at the threshold and overlaid for comparison. **B**) Boxplot showing AP width of PTNs at rheobase from controls and 6-OHDA mice in the presence of BAPTA. **C**) Boxplot showing AP broadening during repetitive firing of PTNs in controls and 6-OHDA mice in the presence of BAPTA. **D**) Frequency-current curve of PTNs in controls and 6-OHDA mice in the presence of BAPTA. **F and G**) Representative AP waveforms at rheobase before and after paxillin application in PTNs from controls (**F**) and 6-OHDA mice (**G**). APs were aligned at the threshold and overlaid for comparison. **H**) Summarized graph showing paxillin broadened APs of PTNs from both controls and 6-OHDA mice. **I**) Summarized graph showing AP broadening during repetitive firing of PTNs from controls and 6-OHDA mice in the presence of paxillin. **J-K**) Representative AP traces of PTNs from controls and 6-OHDA mice in the absence and presence of paxillin. L-M) Frequency-current curves showing paxillin decreased the frequency of PTNs firing om controls, but not in 6-OHDA mice. *, p < 0.05, WSR or MWU tests.

Next, we inhibited BK channel activity by a selective blocker paxillin (10 μM) and found that paxillin significantly decreased fAHP amplitude of PTNs in controls (baseline = 11.2 [7.7, 12.6] mV; paxillin = 8.3 [3.5, 10] mV, n = 12 neurons/3 mice, p = 0.005, WSR), but not in 6-OHDA mice (baseline = 8.7 [7.3, 9.0] mV; paxillin = 8.7 [5.9, 9.8] mV, n = 12 neurons/3 mice, WSR). Consistently paxillin significantly increased AP half-width of PTNs at rheobase in controls (baseline = 0.7 [0.66, 0.79] ms, paxillin = 0.83 [0.78, 0.91] ms, n = 12 neurons/3 mice, p = 0.0024, WSR, Figure 7F, H), but not in 6-OHDA mice (baseline = 0.85 [0.8, 0.97] ms, paxillin = 0.88 [0.83, 0.95] ms, n = 12 neurons/3 mice, p = 0.27, WSR, Figure 7G, H). Thus, blockade of BK channels by paxillin abolished the difference in AP half-width of PTNs at rheobase in control and 6-OHDA mice (AP half-width in paxillin, p = 0.22 between groups, MWU, Figure 7H). These results suggests that BK channel activation contributes to AP repolarization and width of PTNs in controls, but not in 6-OHDA mice.

Further, PTNs in 6-OHDA mice exhibited stronger activity-dependent AP broadening during repetitive firing relative to those from controls (Figure 5D). However, we found that inhibition of BK activity using paxillin abolished difference in the width of the first AP of PTNs (the width of the first AP in paxillin, control = 0.76 [0.71, 0.8] ms, 6-OHDA = 0.87 [0.73, 0.95] ms, p = 0.1, MWU, Figure 7I), but did not affect the robust AP broadening during repetitive firing in 6-OHDA mice relative to that in controls (the width of the last AP in paxillin, control = 0.94 [0.85, 1.06] ms, 6-OHDA = 1.06 [1.0, 1.2] ms, p = 0.01, n = 12 neurons/3 mice for each group, Figure 7I). These results suggest that an unidentified ionic conductance, other than BK channels, also contributes to the activity-dependent AP broadening of PTNs in both controls and 6-OHDA mice, which may play a dominant role in parkinsonian state.

Last, we tested whether BK channel dysfunction contributes to the impaired excitability of PTNs in parkinsonism. In agreement with its role in AP repolarization, blockade of BK channels by paxillin decreased the excitability of PTNs in controls (spike numbers at the maximal current injection, baseline = 55 [49, 59], paxillin = 44 [36, 55], n = 12 neurons/3 mice, p = 0.0015, WSR, Figure 7J, L), but not in 6-OHDA mice (baseline = 37 [20, 47], paxillin = 30 [21, 40], n = 12 neurons/3 mice, p = 0.12, WSR, Figure 7K, M). However, paxillin did not abolish the difference in the firing frequency of PTNs between groups (maximal firing frequency, before paxillin p = 0.0002 between groups, after paxillin, p = 0.0047, MWU), suggesting BK channels dysfunction partially contributes to the decreased intrinsic excitability of PTNs in parkinsonism.

### DA cannot recue the decreased intrinsic excitability of PTNs in parkinsonism

M1 receives dopaminergic inputs from the ventral tegmental area and these neurons are partially degenerate in 6-OHDA mice (Hosp et al., 2011, 2015; Hosp and Luft, 2013). Next, we tested whether a disrupted mesocortical DA neuromodulation contributes to the intrinsic adaptations of PTNs in M1. DA receptors on PTNs were activated through bath perfusion of DA (10 μΜ). However, activation of DA receptors produced negligible effects to the intrinsic excitability of M1 PTNs in controls (e.g., the maximal firing frequency, baseline = 56 [50, 60] Hz; DA = 59 [43, 63] Hz; n = 10 neurons/3 mice; p = 0.9, WSR; Figure. 8A, C, E). In addition, application of DA (10 μM) did not produce significant effects to the intrinsic excitability of M1 PTNs in 6-OHDA mice either (e.g., the maximal firing frequency, baseline = 37 [34, 47] Hz, DA = 39 [35, 45] Hz, n = 12 neurons/3 mice; p = 0.7, WSR; Figure 8B-E). Further, we found that DA (10 μM) did not alter the AP width of PTNs in both groups either (controls: baseline = 0.7 [0.63, 0.74] ms, DA = 0.7 [0.62, 0.73] ms, n = 10 neurons/3 mice, p = 0.77, WSR; 6-OHDA: baseline = 0.78 [0.71, 0.82] ms, 6-OHDA DA = 0.79 [0.67, 0.87] ms, n = 12 neurons/3 mice, p = 0.8, WSR, Figure 8F). These data suggested that restoration of local DA neuromodulation cannot rescue the impaired intrinsic excitability and AP broadening of M1 PTNs following the loss of midbrain DA neurons.

**Figure 8.**
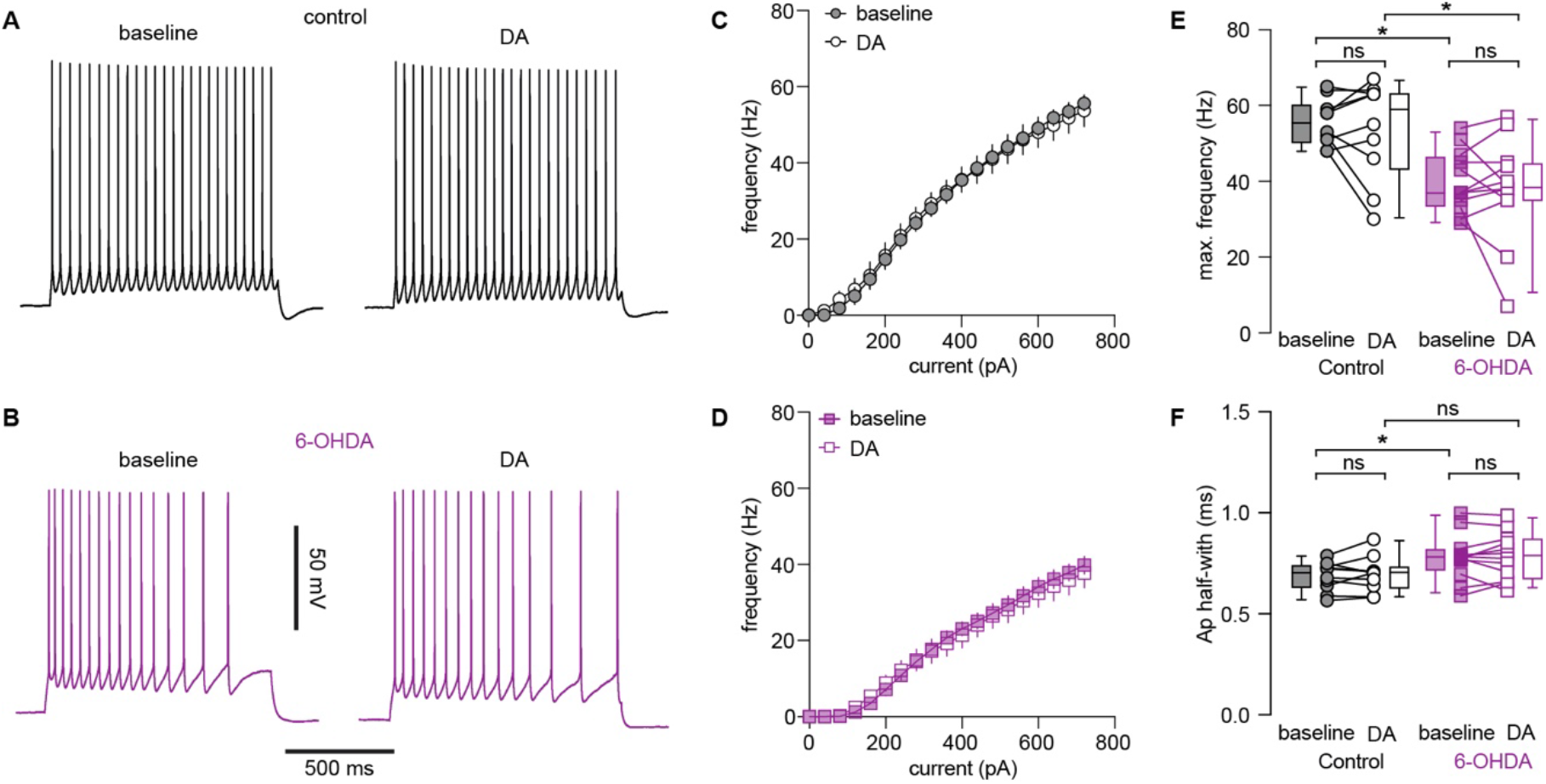
DA application could not rescue the decreased excitability of PTNs following loss of midbrain DA neurons. **A-B**) Representative AP traces of PTNs from controls and 6-OHDA mice in the absence and presence of DA. **C-D**) Frequency-current curves of PTNs from controls (C) and 6-OHDA mice (D) in the absence and presence of DA. **E**) Summarized graphs showing the lack of impact of DA on maximal firing frequency of PTNs from controls and 6-OHDA mice. **F**) Summarized graphs showing the lack of impact of DA on AP width of PTNs from controls and 6-OHDA mice *, p < 0.05, WSR or MWU tests.

## Discussion

We studied changes in the intrinsic properties of M1 pyramidal neurons in experimental parkinsonism. We found a cell-type-specific decrease of the intrinsic excitability of M1 PTNs, but not the ITNs, in parkinsonism. We demonstrated that impaired persistent Na^+^ channel function led to an elevated AP threshold and played a major role in the decreased intrinsic excitability of M1 PTNs. In addition, M1 PTNs also exhibited broadened APs, which partially contributes to the decreased excitability of PTNs during repetitive firing. Further, we revealed that impaired BK channel function underlies the broadened APs of M1 PTNs in parkinsonism. Last, we showed that the decreased intrinsic excitability of M1 PTNs were not reversible by the acute activation of dopaminergic receptors. Our findings strongly suggest that the loss of midbrain DA neurons not only alters basal ganglia activity, but also triggers intrinsic adaptations in M1 circuitry, which contribute to the pathophysiology of parkinsonism.

Cell-type-specific changes in spontaneous activity of M1 pyramidal neurons were first reported in MPTP-treated primates by Pasquereau and Turner (2011). In their excellent work, the authors demonstrated a decreased firing rate and an increased firing irregularity of spontaneous activity of antidromically identified PTNs; in contrast, the firing rate and pattern of corticostriatal neurons were largely intact in MPTP-treated monkeys (Pasquereau and Turner, 2011). Given the different anatomical projections and function of M1 neuron subtypes in motor control, these subtype-specific changes in neuronal activity have great implications in understanding of pathophysiology of parkinsonian motor signs. Yet, few studies have been conducted to understand the cellular and synaptic mechanisms underlying those changes in M1 neuronal activities in parkinsonism. To address this question, we employed *ex vivo* electrophysiology to study the intrinsic properties of projection-defined M1 pyramidal neurons in parkinsonism. Following chronic midbrain DA degeneration, PTNs, but not the ITNs, in the L5b of M1 showed decreased intrinsic excitability and this finding is consistent with the earlier findings in MPTP-treated primates (Pasquereau and Turner, 2011). Given the innervation of subcortical motor centers by M1 PTNs, these results suggested that intrinsic adaptations of M1 pyramidal neurons contribute to an insufficient motor cortical outputs in parkinsonism, in addition to an elevated basal ganglia inhibition as proposed by the classical model (Albin et al., 1989; DeLong, 1990; Galvan and Wichmann, 2008). Thus we propose that these adaptations are important aspects of pathophysiology of parkinsonism.

The decreased excitability of PTNs in 6-OHDA mice associated with a depolarized AP threshold and could be normalized by *I*_Nap_ blockade using low concentration of TTX. These results, together with the disrupted AP rising kinetics of PTNs in 6-OHDA mice, suggest that an impaired *I*_Nap_ function decreases the availability of Na_T_ channels that underlie AP depolarization and further promotes their inactivation during repetitive firing, leading to the decreased excitability of PTNs. We posit that the decreased intrinsic excitability makes neurons are more sensitive to synaptic inputs, as that occurs in basal ganglia neurons in parkinsonism (Wilson 2013). Thus, it is likely that the increased bursting pattern of activity of PTNs *in vivo* can be largely triggered by synaptic inputs, but the enhanced spike adaptation leads to a reduced intraburst frequency in MPTP-treated monkeys (Pasquereau and Turner, 2011; Pasquereau et al., 2016). Further studies will be needed to determine how synaptic inputs shape M1 neuronal activity in parkinsonism.

APs were significantly broadened in PTNs of parkinsonian animals (Figures 4A, 5A, 5B). AP width has significant impact on Ca^2+^ influx during APs (Bean, 2007), yet functional implications of AP broadening in intracellular Ca^2+^ dynamics remain unknown. We showed that the AP broadening in PTNs in parkinsonism was occluded in the presence of BAPTA and paxillin, indicating a key role of BK channels in AP waveform modulation (Shao et al., 1999; Faber and Sah, 2003; Bean, 2007). It has been proposed that inactivation of BK channels could reduce Na_T_ availability for the subsequent AP generation during repetitive firing, leading to a decreased neuronal excitability (Bean, 2007; Jaffe et al., 2011). Consistently, BK inhibition by paxillin strongly decreased excitability of PTNs in controls, but not in 6-OHDA mice, leading to a partial abolishment of the difference in neuronal excitability between groups (Figure 7L, 7M). Thus, we propose that impairments of both *I*_Nap_ and BK activities exacerbate progressive accumulation of Na_T_ inactivation during repetitive firing, leading to the decreased intrinsic excitability of PTNs in parkinsonism (Fleidervish et al., 1996). It is worthy to note that PTNs in both controls and 6-OHDA mice exhibited activity-dependent AP broadening in the presence of BK inhibition (Figures 7C, 7I). It indicates that unidentified ionic channels also contribute to AP repolarization of PTNs (Rudy and McBain, 2001; Pathak et al., 2016; Soares et al., 2017), and their inactivation underlies broadened APs in both groups, but possibly plays a major role in AP repolarization in parkinsonism.

An acute application of DA did not affect the excitability of PTNs in both controls and in 6-OHDA mice. These results are consistent with recent findings that dopaminergic modulation of pyramidal neurons is mainly mediated indirectly by parvalbumin-expressing interneurons in M1 (Vitrac et al., 2014; Cousineau et al., 2020). Alternatively, it is possible that loss of mesocortical dopaminergic modulation contributes to the altered intrinsic excitability of PTNs, but such changes are not reversible by acute dopaminergic receptor activation. Compelling evidence suggests a critical role of dopaminergic modulation in the structural and functional plasticity of the motor cortical circuits (Lindenbach and Bishop, 2013; Ueno et al., 2014; Guo et al., 2015b). Last, circuitry mechanisms may also underlie cellular adaptions in M1 in parkinsonism. For example, recent studies highlighted a critical role of indirect pathway striatal projection neurons in driving cellular and synaptic adaptations in basal ganglia nuclei in parkinsonism, such as the subthalamic nucleus (Chu et al., 2017; Sharott et al., 2017; Ryan et al., 2018; McIver et al., 2019). Similarly, pathological basal ganglia outputs in parkinsonism can drive adaptive changes in M1 via the thalamocortical projections (Villalba et al., 2021).

The exaggerated rhythmic activity and aberrant oscillations in the basal ganglia circuitry are closely related to the expression of motor symptoms in PD (Bevan et al., 2002; Little and Brown, 2014; Hemptinne et al., 2015). Indeed, the motor cortex exhibits abnormal neural activity in parkinsonian state at both the single-cell and population levels (Goldberg et al., 2002; Pasquereau and Turner, 2011; Hemptinne et al., 2015; Pasquereau et al., 2016). Both experimental and computation studies have indicated that cerebral cortical inputs play a crucial role in orchestrating pathological activity in basal ganglia nuclei (e.g., the external global pallidus and the subthalamic nucleus) (Bevan et al., 2002; Magill et al., 2004; Kita and Kita, 2011; Corbit et al., 2016; Chu et al., 2017; Sharott et al., 2017). Recent studies also showed that electrical or optogenetic stimulation of the motor cortical pyramidal neurons and/or their projections produced therapeutic effects in parkinsonian animals (Gradinaru et al., 2009; Li et al., 2012; Sanders and Jaeger, 2016; Magno et al., 2019). Thus, an in-depth understanding of motor cortical adaptations in PD at the cellular and circuitry levels will help to design better therapeutic approaches with fewer off-target effects.

## Acknowledgements

This work was supported by NINDS grant R01NS121371 and the startup fund from Van Andel Institute (H.Y.C). The authors thank Mr. David Nadziejka for technical editing.

## Notes

**Conflict of interest**: The authors declare no competing financial interests.

### Competing Interest Statement

The authors have declared no competing interest.

### Summary of Updates

new results were added and texts were revised.

